# Identification of modulated whole-brain dynamical models from nonstationary electrophysiological data

**DOI:** 10.1101/2025.05.01.651677

**Authors:** Addison Schwamb, Zongxi Yu, ShiNung Ching

## Abstract

**Objective:** Understanding the mechanisms underlying brain dynamics is a long-held goal in neuroscience. However, these dynamics are both individualized and nonstationary, making modeling challenging. Here, we present a data-driven approach to modeling nonstationary dynamics based on principles of neuromodulation, at the level of individual subjects.

**Approach:** Previously, we developed the mesoscale individualized neural dynamics (MINDy) modeling approach to capture individualized brain dynamics which do not change over time. Here, we extend the MINDy approach by adding a modulatory component which is multiplied by a set of baseline, stationary connectivity weights. We validate this model on both synthetic data and publicly available EEG data in the context of anesthesia, a known modulator of neural dynamics.

**Main Results:** We find that our modulated MINDy approach is accurate, individualized, and reliable. Additionally, we find that our models yield biologically interpretable inferences regarding the effects of propofol anesthesia on mesoscale cortical networks, consistent with previous literature on the neuromodulatory effects of propofol.

**Significance:** Ultimately, our data-driven modeling approach is reliable and scalable, and provides insight into mechanisms underlying observed brain dynamics. Our modeling methodology can be used to infer insights about modulation dynamics in the brain in a number of different contexts.

## 1. Introduction

An important and persistent challenge in the analysis of recorded mesoscale neural activity (i.e., commensurate with externally recorded fields and potentials) is the inference of latent neurophysiological mechanisms that underlie overt observations. Indeed, while there are myriad tools and methods to analyze brain electrophysiological recordings (e.g., power spectral density estimation [1]), there are fewer that provide direct inference of circuit-level mechanisms (i.e., the interaction of excitatory and inhibitory neural subpopulations, from which said recordings originate). Identifying and understanding these mechanisms is a difficult task, however, as they are not directly observable via typical mesoscale recording modalities. For instance, electroencephalography (EEG) measures electrical potential non-invasively at the scalp, and hence does not provide direct access to neuronal sub-population-level activity [2]. Parametric dynamical systems modeling offers a methodological path to obviating this issue. Such models, via their mathematical formulation, embed mechanistic hypotheses or inductive biases regarding how neural activity and secondary observables such as EEG are generated. While dynamical systems modeling is potentially powerful in this context, there are several extant challenges that remain unsolved regarding the construction of such models from neural recordings.

First, neural activity patterns vary between individuals, indicating that the underlying mechanisms also vary on an individual level. For example, the posterior dominant rhythm is a classic example of neural dynamics, which manifests as a strong alpha-band (i.e., 10-14 Hz) oscillation localized to the posterior of the scalp in EEG recordings (i.e., overlaying occipital cortex). While the general characteristics of the posterior dominant rhythm (its frequency and spatial location) are consistent across individuals [3], the exact frequency and power of the posterior dominant rhythm is specific to individuals [4]. To address this challenge of individuality, work has been done to create data-driven models based on single-subject data [5, 6], providing individualized neural dynamics models which can be used to infer mechanisms underlying a person’s brain activity [7, 8, 9]. This approach is schematized in Figure 1a.

**Figure 1.**
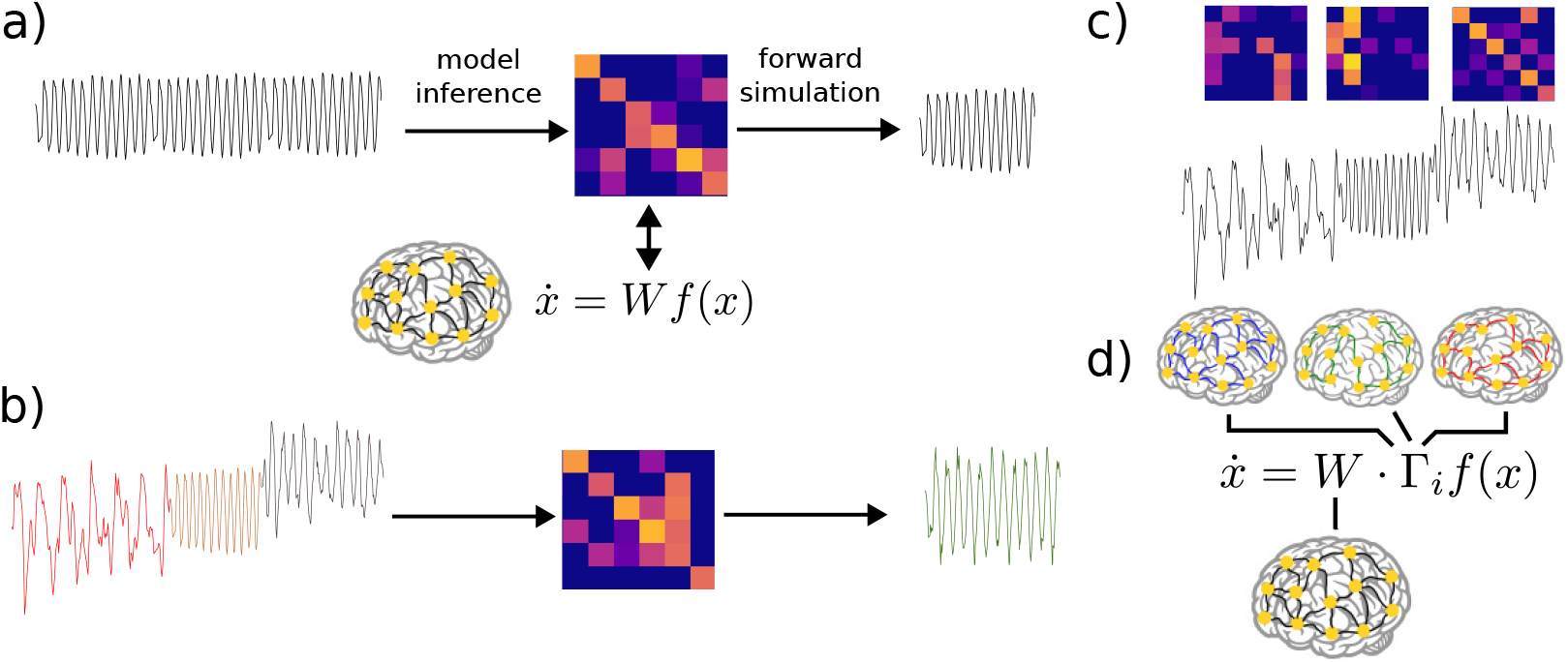
Approaches to fitting individualized models from data. a) In a stationary setting, a single set of (time-invariant) model parameters (here, a connectivity/weight matrix) is fit to data, yielding a corresponding stationary prediction/forward simulation. b) When data is non-stationary, fitting a single stationary model will in general lead to erroneous or averaged modeled dynamics. c) A typical way to address non-stationary data is to fit separate models (e.g., distinct weight matrices) to the distinct non-stationary regimes. d) In the proposed approach, we seek to fit non-stationary models that decompose parameters into a baseline connectivity and regime-specific modulatory matrices.

In addition to varying between individuals, neural dynamics are also highly non-stationary, i.e., they vary temporally based on many factors such as modulation of neurophysiologic states (e.g., sleep vs. wake [10], rest vs. task [11], and healthy activity vs. pathology [12]). Therefore, an individual dynamical systems model of the kind mentioned above can typically only offer insights for a specific physiological regime, and/or for a relatively narrow epoch of time during which dynamics are stationary [13]. There is an unmet need for data-driven modeling methods that can capture non-stationarity in dynamics associated with multiple neurophysiologic states. If an approach is taken that does not account for this nonstationarity, the model which is returned will be a poor fit for any of the regimes present in the data (Figure 1b).

To overcome this challenge, several approaches to time- or state-dependent dynamical systems models for neural activity have been suggested. In essence, these models embed a mechanism to modify the model parameters in a manner that captures categorical changes in neural dynamics (e.g., sleep vs. wake). For instance, the switching linear dynamical system (SLDS) framework [14, 15] embeds multiple linear dynamical systems, of which a single model is “active” at any given time [16, 17, 18], as shown in Figure 1c. Switches between models are often enacted through a latent model, such as a hidden Markov model [19] that provides the dynamics behind switches [20, 21]. Through such a mechanism, a model can embed a number of distinct dynamical regimes. A similar approach is taken in [22, 23], but here the researchers use a recurrent neural network (RNN) model, rather than a linear dynamical system. They construct a discrete number of RNNs (each with a different recurrent weight matrix), thus implementing a different set of dynamics for each RNN.

It is important to note that modeling non-stationarity can be understood as a problem with two phases: i) modeling when dynamical regimes change, and ii) modeling what in the latent dynamics has changed. Our proposed approach tackles the latter phase. Specifically, we infer the changes in latent dynamics as changes in mesoscale *neuromodulation* (Figure 1d), rather than comparing distinct dynamic regimes (Figure 1c). In this paper, we take a similar approach to [22, 23], beginning with a biologically interpretable RNN model describing population-level neural activity. However, in contrast to the approach of having a discrete set of recurrent weight matrices, we implement a modulation architecture (Figure 1d). To our knowledge, data-driven inference of neuromodulatory effects on mesoscale dynamics has not been previously pursued in this way.

Our model consists of a single weight matrix common to all dynamical regimes, which is then multiplied element-wise by one of a discrete number of modulatory matrices corresponding to different dynamical regimes. Such an assumption is based, schematically, on the actions of neuromodulators on neural circuits that modulate synaptic efficacy. We assume that such modulation occurs on a slower timescale than the timescale of neural activity. By using this modulation structure, we are able to separate the aspects of an individual’s dynamics which are common across time and neurophysiologic state from those which are variable. Additionally, we impose specific excitatory and inhibitory sub-structures on our parameters, in order to preserve the biological interpretability of our fitted models. All model parameters are estimated directly from data in an individualized manner.

Below, we develop and present the proposed methodology. We first introduce our modulated recurrent neural network architecture, building from our prior data-driven modeling approach of Mesoscale Individualized Neural Dynamics (MINDy) [8, 9] to enact the aforementioned modulation architecture. We then validate this on simulated data, to test the accuracy and reliability of our model in recovering known ground truth parameters. We then test our model on the ability to infer distinct neural mechanisms associated with levels of general anesthesia, a pharmacological modulation of neural circuits. These data allow us to further validate the reliability and individuality of our fitted models. We then reconcile the inferred models and mechanisms with prior observations regarding the modulatory effects of anesthesia on cortical networks.

## 2. Methods

### 2.1. Model formation

Our model is adapted from our prior whole-brain dynamical systems framework in [8, 9]. The base (i.e., unmodulated) dynamics are governed by:

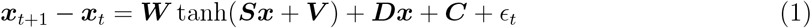

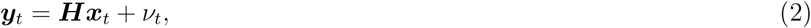

where ***x***_*t*_ ∈ ℝ^*n*^ represents the neural activity of *n* neural populations at time *t*, and {***W***, ***S, V***, ***D, C***} are the tunable model parameters. Here, **W** is the connectivity matrix, **D** is a diagonal matrix representing the decay (leak) in neural activity for each population, **S** and **V** parameterize the slope and offset of the sigmoidal activation function, and **C** represents a baseline bias of each neural population. ***H*** is the lead field model which translates the population-level neural activity into the measured data, and *ϵ*_*t*_ and *ν*_*t*_ represent the process noise and measurement noise, respectively.

Note that this model is mathematically comparable to a vanilla recurrent neural network, but arranged and constrained to reflect biologically interpretable relationships between excitatory and inhibitory neural populations. By assigning neural populations to spatial locations in the brain and constraining the parameters to specific valence (i.e., positive/excitatory, negative/inhibitory), we can examine changes in these parameters associated with physiological changes. Specifically, we enforce excitatory and inhibitory substructures onto the neural population activity vector ***x***_*t*_ and the connectivity matrix ***W*** like so:

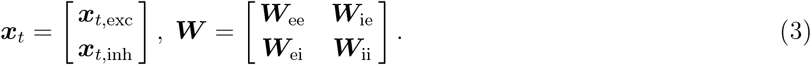

Entries in ***W***_ee_ and ***W***_ei_ are constrained to be positive or 0, as they represent the connections from excitatory neural populations. Additionally, ***W***_ee_ and ***W***_ei_ are full submatrices. Conversely, ***W***_ie_ and ***W***_ii_ have entries which are negative or 0, as they represent the connections from inhibitory neural populations. Since inhibitory neurons only have local connections [24], these submatrices are constrained to be diagonal.

We adapted and generalized this model by adding neuromodulation via matrices ***Γ***_*i*_, *i* = 1, …, *m* multiplied element-wise by the connectivity matrix ***W*** :

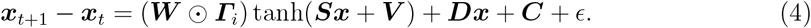

Here, *m* represents the number of modulatory states to be modeled. This modulation structure is schematized in Figure 1d. To preserve the signed connectivity structure of ***W***, entries in each ***Γ***_*i*_ matrix are constrained to be positive or 0. This enables a modulation which scales entries in ***W*** by varying amounts, without affecting the base structure of the connectivity.

The specification of ***H*** is made to reflect the specifics of the data modality being used for model construction. In our case, because we are using cortical data (EEG), we account for prevailing hypotheses regarding the contributions of different cell types to surface-level potentials. In this context, it is generally believed that inhibitory neurons are not close enough to the surface of the cortex, nor do they possess the spatial organization, to generate fields detectable via EEG [25]. Therefore, we construct our lead field matrix ***H*** to have zeros in the submatrix multiplied by the inhibitory component of ***x***:

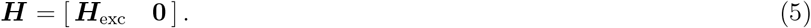

### 2.2. Model fitting procedure

Because ***H*** is not invertible, we now face what is sometimes termed a dual inference problem: i) estimate ***x***_*t*_ and ii) estimate the parameters {***W***, ***Γ***, ***D, S, V***, ***C***}, from the observable (i.e., EEG) data.

To address this, we adopt the iterative estimation approach detailed in [9, 26]: first we apply a Kalman filter [27, 28] to a small window of data in order to estimate the state. We then evolve the model forward from the (estimated) state to generate a forward prediction. We then backpropagate the error through both the free simulation steps and the Kalman filtering steps, calculating the error gradient for each parameter at each step. In addition to calculating the parameter error gradients (i.e., error gradients of {***W***, ***Γ***, ***D, C, S, V*** }), we also calculate the error gradients of the estimated covariances of the noise terms *ϵ* and *ν*. These noise covariances are necessary for the Kalman filter to estimate the current state ***x***_*t*_, and are optimized alongside the model parameters. By backpropagating the error in this manner and fitting both the parameters and the noise covariances, we fit a model which produces the most accurate state estimator which can best predict future measurements.

After backpropagation through the free simulation and Kalman filtering steps, we then update the model parameters and the estimates of the noise covariances. Then, the process is repeated with a new small window in the epoch of fitting data. This process of Kalman filtering, free simulation and backpropagation is repeated with new data windows until the Kalman and free simulation errors converge. Multiple windows of the data epoch are used so that the model captures the general dynamics and statistical properties of the entire epoch, to avoid overfitting to a few timesteps of data. These windows are selected at random to avoid biasing the fit toward any one period within the epoch (e.g., the beginning or end of the epoch).

In the unmodulated MINDy model, the parameters do not vary temporally, so all parameters are updated in all steps of the model. In the expanded modulated MINDy model proposed here, only one of the ***Γ***_*i*_ matrices is used at each timepoint. To account for this, we select the appropriate ***Γ***_*i*_ at each time *t* using the modulation labels and use that ***Γ***_*i*_ for model evolution and gradient calculation.

To aid in the tractability of the problem, we implement several constraints on ***W*** and ***Γ***. We constrain ***W***_*ee*_ and ***W***_*ei*_ to have 75% of their non-diagonal connections be zero, enforcing a prior level of sparsity in connections. We also constrain each ***Γ***_*i*_ to be rank 1, i.e., the outer product of two *n*-dimensional vectors. This assumption is motivated by the premise that neuromodulators act in a spatially diffuse manner [29]. Because each ***Γ***_*i*_ is multiplied element-wise by ***W***, the excitatory submatrices of each ***Γ***_*i*_ also effectively have 75% of their elements set equal to 0.

To fit the model parameters, we use NADAM [30], implemented with PyTorch’s Autograd engines [31]. This improves fitting efficiency and scalability relative to our prior work, by allowing GPU acceleration, which can be especially beneficial for processing large-scale EEG recordings. Notably, this significantly reduces the number of iterations needed for the convergence of the backpropagated Kalman filter approach. With Autograd, both the covariance update and local linearization process are included in the backward operation.

### 2.3. Simulation and actual data

#### 2.3.1. Synthetic data

To enable the validatation of our fitting procedure, we created synthetic data with known parameters. To create this synthetic data, we established models with a combination of fixed and random parameters. The fixed values and distributions for the model parameters are listed in Table 1. It should be noted that our ***W*** submatrices were constructed as a linear combination of a sparse and low-rank matrix via:

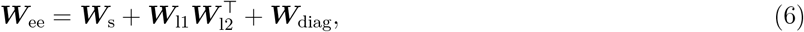

where ***W***_s_ ∈ ℝ^*n*×*n*^ is a sparse matrix, ***W***_l1_, ***W***_l2_ ∈ ℝ^*n*×*n/*4^ are low rank matrices, and ***W***_diag_ ∈ ℝ_*n*_ is a vector specifying the diagonal self-connection weights. Note that (6) specifies ***W***_ee_, but both ***W***_ee_ and ***W***_ei_ were constructed in this way. ***W***_ie_ and ***W***_ii_ were constructed as diagonal matrices with diagonal values directly sampled from their distributions.

**Table 1.**
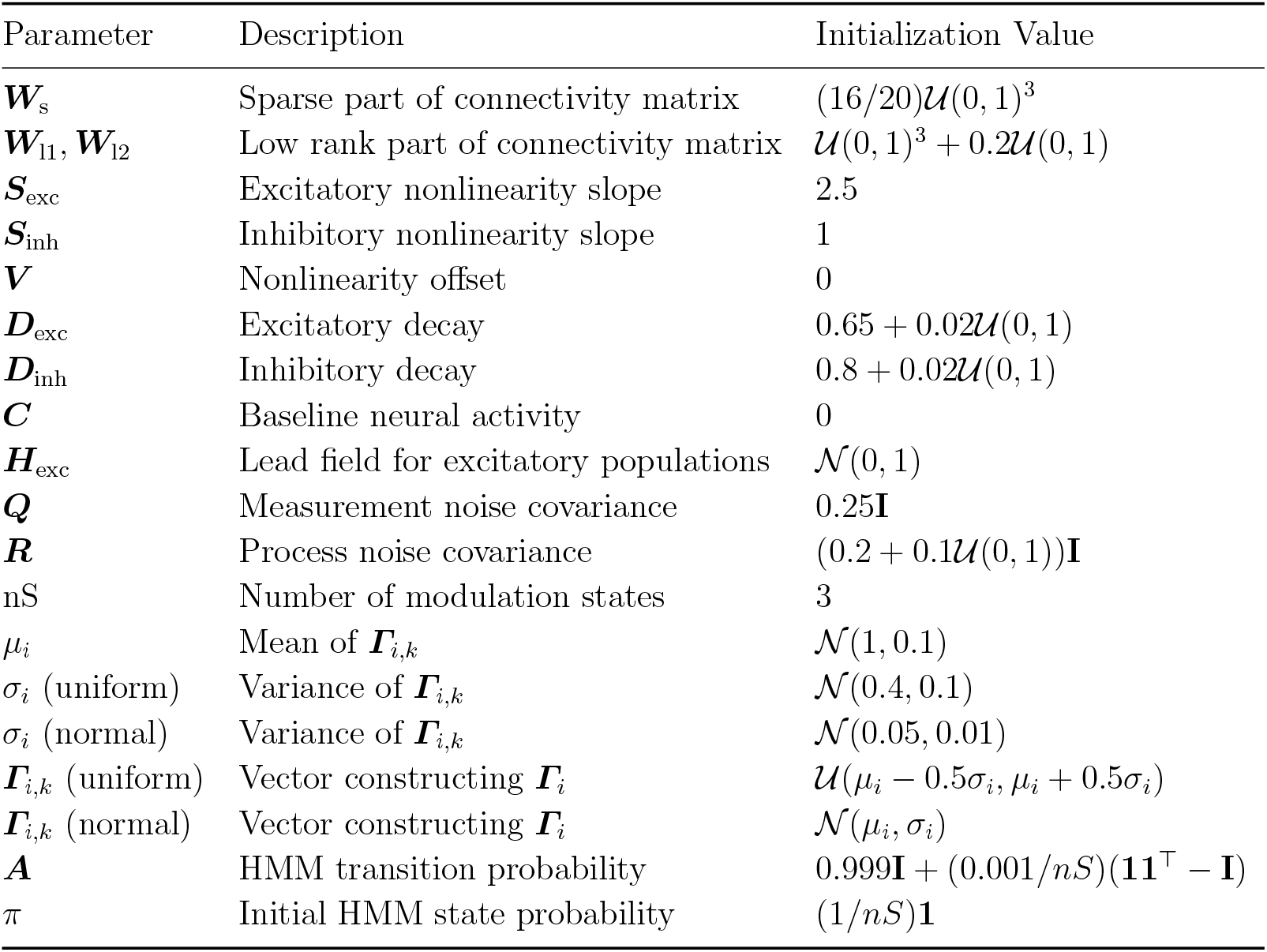
Parameter values or random distributions used to create synthetic data models. **I** denotes the identity matrix, and **1** denotes the vector of all ones.

We initialized each ***Γ***_*i*_ matrix as the outer product of a vector drawn from a unique distribution. To construct each unique distribution, we first randomly generate its mean, *µ*_*i*_, from 𝒩 (1, 0.1). We then generate a random binary digit indicating whether we should use a uniform distribution, or a normal distribution. Then, we generate a variance *σ*_*i*_ for the distribution. If we are using a uniform distribution, *σ*_*i*_ ~ 𝒩 (0.4, 0.1), and if we are using a normal distribution, *σ*_*i*_ ~ 𝒩 (0.05, 0.01). Then, we generate a vector, **Γ**_*i*,*k*_, from the constructed distribution: if uniform, **Γ**_*i*,*k*_ ~ 𝒰(*µ*_*i*_ − (*σ*_*i*_*/*2), *µ*_*i*_ + (*σ*_*i*_*/*2)); if normal, **Γ**_*i*,*k*_ ~ 𝒩 (*µ*_*i*_, *σ*_*i*_). Then, 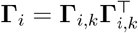.

We chose the distributions for *µ*_*i*_ and *σ*_*i*_ such that there could be variation in the distributions of **Γ**_*i*_, while also maintaining values close to 1. If the values of **Γ** are very large or very small, **Γ** will overwhelm **W** in the effective neural connectivity **W** ⊙ **Γ**_*i*_. In other words, our assumed modulation does not re-scale synaptic weights by large amounts.

To generate state changes in our synthetic data, we initialized a hidden Markov model (HMM) with transition probability matrix ***A*** and initial state probability *π*. At each timestep, we calculated the probability of the next state based on ***A*** and then randomly selected the state index weighted by the calculated probability.

Once we had generated our random models, we forward simulated them for 20,000 timesteps (equivalent to 80s of 250Hz EEG) to create synthetic data. When fitting on this synthetic data, the true lead field matrix and noise covariances (***H, Q***, and ***R***) were used to initialize the models, but the other model parameters were initialized randomly and compared to the true values after fitting, to test parameter recovery when ground truth is known. Since the true value of ***W*** is known, however, we create a mask zeroing out the same entries in the fit ***W***, to avoid zeroing out connections which are actually present in the synthetic models.

#### 2.3.2. EEG data

For our second experiment, we used EEG data of 20 subjects dosed with propofol published in [32] and available online [33]. In this data, subjects were recorded at four levels of sedation: i) resting baseline with no anesthesia, ii) mild sedation (defined as 0.6 *µ*g/mL target blood plasma concentration), iii) moderate sedation (1.2 *µ*g/mL target blood plasma concentration), and iv) recovery from sedation. Each sedation level had approximately 7 minutes of data, recorded after 10 minutes of allowing the blood plasma level to reach a steady state and was saved as a separate EEG file. This data is highly compatible for our purposes because the four pharmacological regimes above can be used as labels for constructing and validating our modulated model.

We filtered the data between 0.5 and 15Hz, subtracted the median of each channel, and divided by the mean absolute deviation of each channel. We combined all four sedation state recordings into a single array for each subject, and added a sedation state index *i* ∈ {1, 2, 3, 4} for each timepoint of the array. We also used a subset of the recorded channels in [32], using only 20 channels, shown in Figure 2.

**Figure 2.**
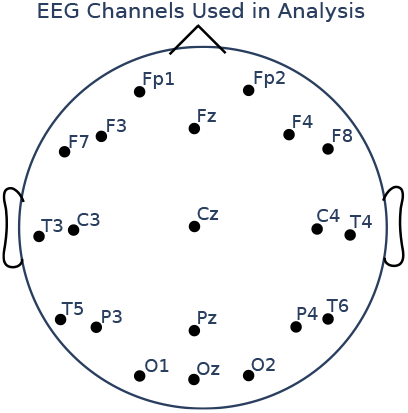
EEG channels used in analysis of propofol dosage EEG data.

Our ***H*** matrix was constructed as in (5), with

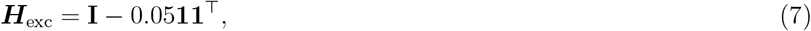

where **I** denotes the 20×20 identity matrix and **1** denotes the 20-dimensional vector of all ones. We constructed ***R*** and ***Q*** as diagonal matrices, with ***R*** = 1.2**I** and ***Q*** = 0.25**I**. The other model parameters were initialized randomly, as in experiment 1. In this experiment, we do not have a ground truth of the 75% of non-diagonal connections in the ***W***_ee_ and ***W***_ei_ submatrices which are zero as we did in the synthetic data case. To continue enforcing this constraint, we selected a random 75% of non-diagonal connections in ***W***_ee_ and ***W***_ei_, and set these to 0 for all subjects. Finally, because we wanted to explore modulations relative to baseline (prior to propofol dosing), we enforce ***Γ***_1_ = **11**^⊤^, i.e., ***W*** ⊙ ***Γ***_1_ = ***W***. The other ***Γ*** matrices (*i* ∈ {2, 3, 4}) are fit as all ***Γ***_*i*_ in the synthetic data experiment.

## 3. Results

### 3.1. Modulated MINDy recovers ground truth connections and modulations

Our first experiment tested whether modulated MINDy could accurately recover the connectivity and modulation matrices in models with known ground-truth parameters. To benchmark this, we compared the accuracy of parameter inferences for the modulated MINDy architecture in the presence of multiple non-stationary regimes to the performance of the unmodulated MINDy architecture (previously validated in [9]) on data in which there is only a single, stationary regime. That is, for the unmodulated MINDy results, we fit unmodulated MINDy models to single-regime synthetic data generated by using the simulated data setup specified in [9].

As shown in Figure 3a, we achieve ground truth correlations of nearly the same level as in the unmodulated MINDy problem. The unmodulated MINDy models achieve a median correlation coefficient of *r* = 0.9771 (IQR: 0.9720-0.9866) for the full ***W*** correlation, whereas the modulated MINDy models yield a correlation coefficient of *r* = 0.9280 (IQR: 0.8539-0.9362). The EE and EI submatrices had similar correlations: MINDy EE, *r* = 0.9873 (IQR: 0.9519-0.9957); Modulated MINDy EE, *r* = 0.9202 (IQR: 0.8238-0.9408); MINDy EI, *r* = 0.9715 (IQR: 0.9506-0.9895), Modulated MINDy EI, *r* = 0.9227 (IQR: 0.8615-0.9495). Since we are fitting not only one connectivity matrix but also several ***Γ*** matrices, a slight decrease in fit quality is expected relative to the unmodulated problem.

**Figure 3.**
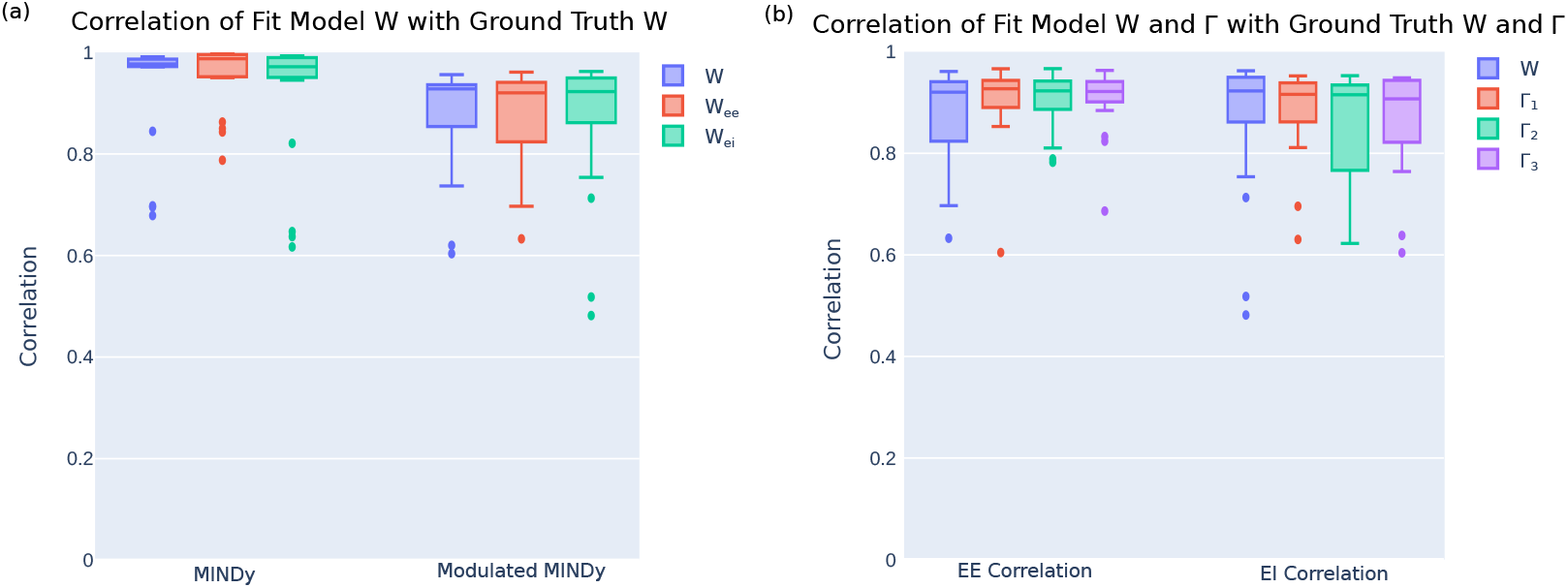
Modulated MINDy returns models with high ground truth correlations. (a) Ground truth correlations for the full connectivity, excitatory-excitatory connections, and excitatory-inhibitory connections for both unmodulated and modulated MINDy problems. (b) Ground truth correlations for the connectivity matrix (***W***) and the connectivity modulation matrices (***Γ***) in the modulated MINDy problem.

Additionally, we achieved similarly high ground truth correlations for the EE and EI components of the modulation matrices ***Γ*** : ***Γ***_1_ *r* = 0.9269 (IQR: 0.8902-0.9434) EE, *r* = 0.9160 (IQR: 0.8618-0.9385) EI; ***Γ***_2_ *r* = 0.9228 (IQR: 0.8864-0.9421) EE, *r* = 0.9152 (IQR: 0.7665-0.9343) EI; ***Γ***_3_ *r* = 0.9215 (IQR: 0.9011-0.9411) EE, *r* = 0.9071 (IQR: 0.8216-0.9435) EI, as shown in Figure 3b. These results indicate that we can accurately recover both the true base connectivity and the true modulations of that connectivity.

Figure 4 provides examples of individual connection and modulation matrices and their recovered estimates via modulated MINDy (i.e., the proposed method). As shown, the spatial patterns of high- and low-amplitude connections in the true matrices are replicated in the estimated matrices. The estimated ***W*** tends to have a slightly higher amplitude than the true ***W***, and the estimated ***Γ*** tends to have a slightly lower amplitude than the true ***Γ***. This is to be expected, since we only regularized the structure of ***W*** and ***Γ***, and did not regularize their amplitude. In summary, we can conclude that the modulated MINDy model can estimate the connectivity and modulation parameters to within a potential scaling factor of the true value.

**Figure 4.**
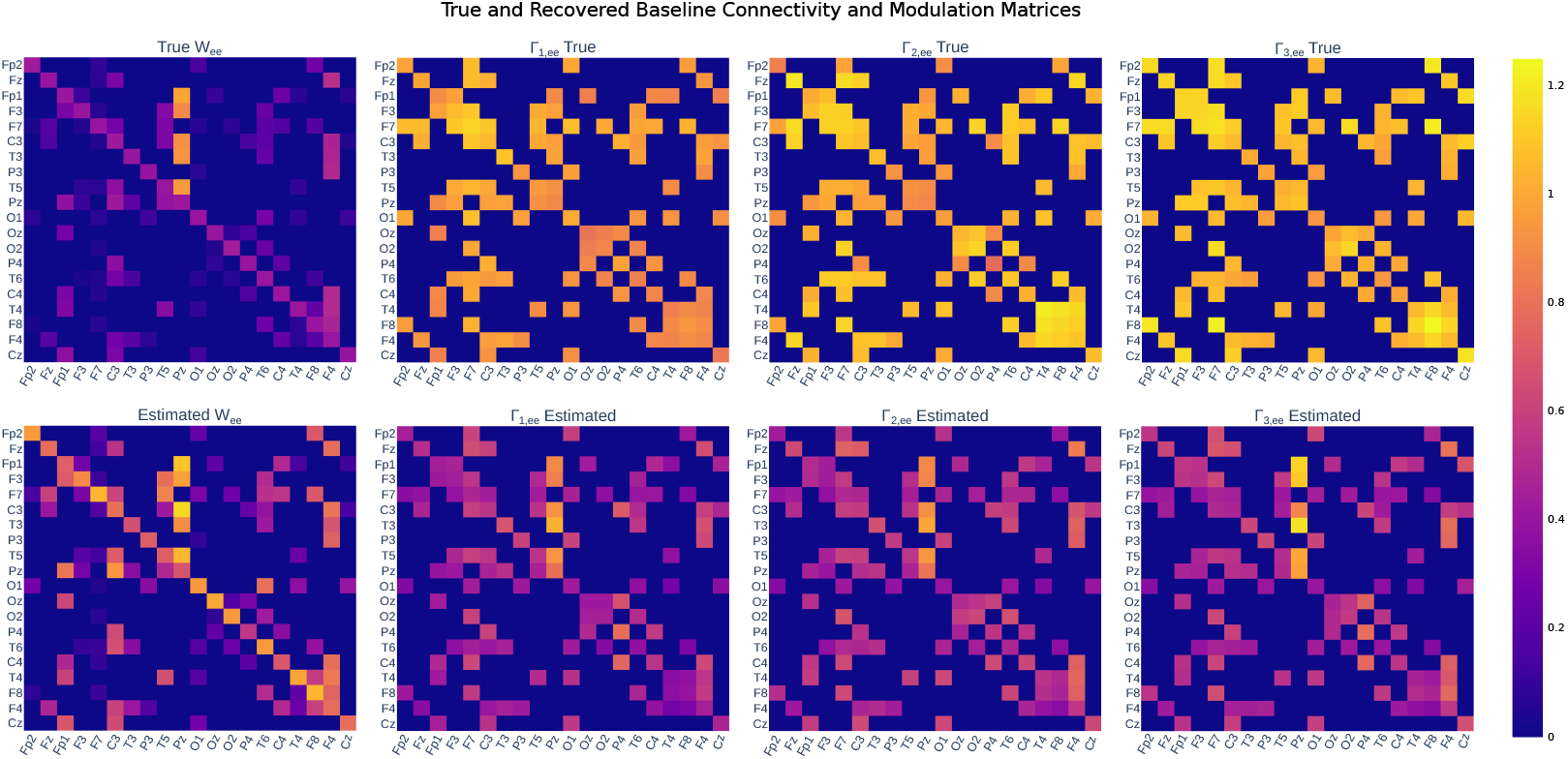
Modulated MINDy returns connectivity and modulation matrices accurate to a potential scaling factor. True (top row) and estimated (bottom row) ***W*** and ***Γ*** matrices from a single subject.

### 3.2. Modulated MINDy is reliable and individualized

In our second experiment, we tested modulated MINDy on actual EEG data to understand how it functioned on a population of individuals. We split each subject’s data in half temporally within each sedation level, in order to perform a test-retest reliability analysis.

We observed that models were reliable, showing high correlations within a single subject (***W*** : *r* = 0.7670 (IQR: 0.7187-0.8118); ***W***_ee_: *r* = 0.7930 (IQR: 0.7276-0.8746); ***W***_ei_: *r* = 0.6855 (IQR: 0.6054-0.7202; Figure 5). Additionally, correlations across subjects were significantly lower (***W*** : *r* = 0.6230 (IQR: 0.5722-0.6802), *p* = 1.97e^−11^; ***W***_ee_: *r* = 0.6510 (IQR: 0.5760-0.7252), *p* = 1.04e^−8^; ***W***_ei_: *r* = 0.5144 (IQR: 0.4275-0.5821), *p* = 1.61e^−8^), indicating that models are individualized, a requisite condition for capturing dynamics and mechanisms which vary across individuals.

**Figure 5.**
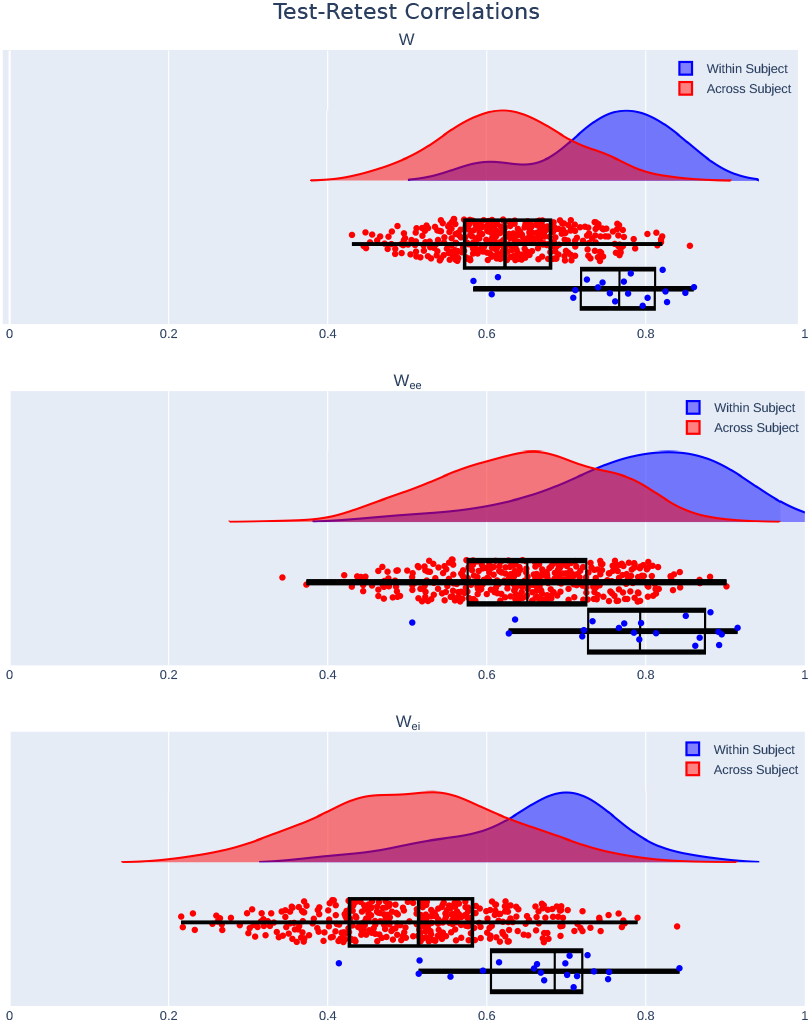
Modulated MINDy is reliable and individualized. Test-retest correlations on split-half data are significantly higher within subject than across subject in the full ***W*** connectivity matrix, as well as the EE and EI submatrices.

### 3.3. Models identify consistent modulation associated with pharmacology

Having established validity in ground-truth settings, we proceeded to apply the method to labeled EEG data from stages of propofol anesthesia. Specifically, for each of 20 individuals, we inferred a base connectivity matrix for the pre-sedation regime, as well as modulation matrices for the regimes of mild anesthesia, moderate anesthesia and recovery. To test our method’s ability to provide insight into the observed changes into dynamics, we analyzed the modulation (***Γ***) matrices at each sedation level compared to baseline.

We observed first that the vast majority of the values within the matrices fall within the range [0, 1), though a small percentage (***Γ***_2_: 0.169%, ***Γ***_3_: 0.150%, ***Γ***_4_: 0.128%) are greater than 1 (Figure 6). Since these values are multiplying the base connection weights ***W***, a fractional value in ***Γ*** indicates a relative decrease of connection strength, or a relative increase in inhibition. In the population average, all values are fractional. This pattern of the modulations increasing inhibition is consistent with the widespread inhibitory effects of propofol [34, 35].

**Figure 6.**
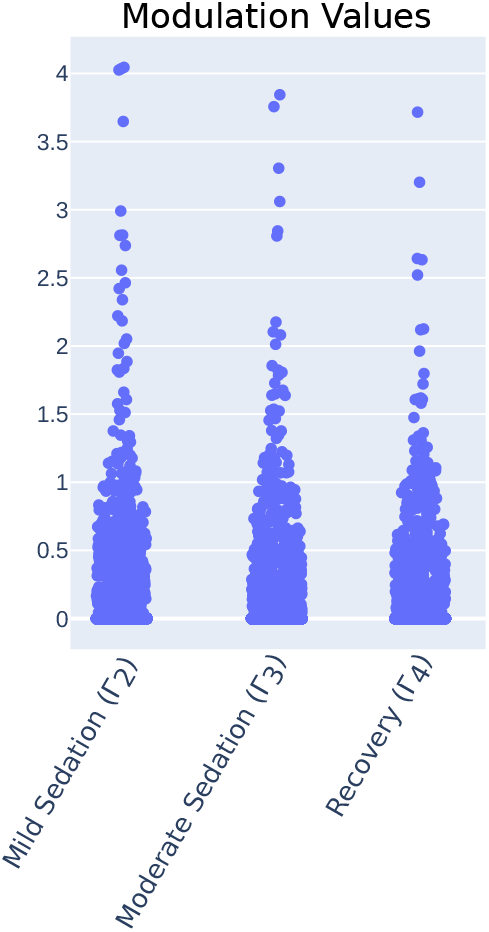
Modulated MINDy identifies inhibitory modulation during sedation. Distributions of all modulation values in each sedation state.

Importantly, we went beyond these bulk inferences and also analyzed the spatiotemporal changes associated with each modulation regime. We found that in the population mean, the modulation values which were highest (i.e., smallest relative inhibition) were typically connections to populations underlying posterior channels, particularly in the moderate sedation state (Figure 7a). This is compatible with prior characterizations of the effect of propofol on attenuating the posterior regions of the scalp [36]. We also noted that each modulation state introduced specific spatial changes to the effective connectivity. To quantify this, we define a measure of the modulation’s impact, defined as the difference between two temporally adjacent modulations matrices scaled by ***W***, e.g., ***W*** ⊙ (***Γ***_2_ − ***Γ***_1_). Thus, we account for both the magnitude differences in ***Γ*** as well as the base magnitude of the connection weights which are being modulated. A positive impact indicates a strengthened connection, and a negative impact indicates a weakened connection.

**Figure 7.**
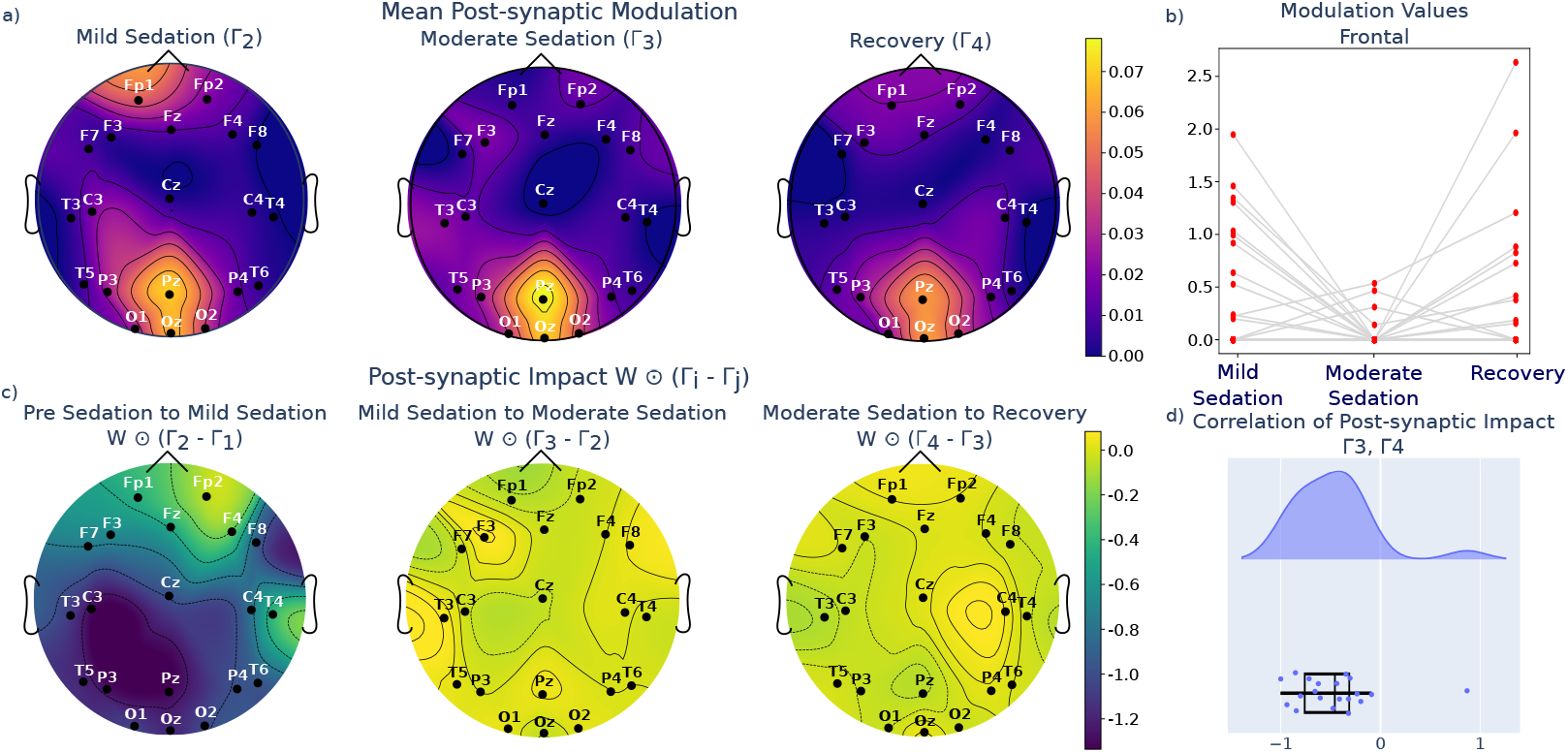
Modulated MINDy identifies inhibitory, spatially focused modulation during sedation. a) Mean post-synaptic modulation in population mean of modulation matrices in sedation states. b) Post-synaptic modulation impact (difference between two temporally adjacent modulation matrices, scaled by ***W***) across transitions between sedation states (population mean). c) Distributions of EE modulation values in frontal channels. d) Correlation between moderate sedation post-synaptic impact and recovery post-synaptic impact.

As the subjects move from pre-sedation to mild sedation, there is a widespread weakening of connections across the whole brain, but again particularly in the posterior areas (Figure 7c, panel 1). As the subjects progress from mild to moderate sedation and from moderate sedation to recovery, the changes are of smaller, less significant magnitude (Figure 7c). The change from mild to moderate sedation is characterized by a further weakening of the frontal connections (Figure 7b). This general observation is consistent with the observation of anteriorization of neural activity in the time before loss of consciousness[36, 37], in which the loss of posterior EEG power is accompanied by a potentiation of activity in frontal regions (note the posterior-anterior dichotomy in inferrered modulatory impact, especially from baseline to mild sedation, panel 1 of Figure 7c). Anteriorization has been modeled as a differential effect of inhibition along the posterior-anterior axis [38, 39]. Interestingly, as the subjects transition from moderate sedation to recovery, there is a reversal of the effects seen in the moderate sedation (Figure 7c), highlighted by the negative correlations between the impact of moderate sedation and the impact of recovery (Figure 7d). This indicates a gradual decrease in EE connection strength as the subjects are dosed with anesthesia, followed by a return to a state similar to the mild sedation in the recovery phase.

Overall, these results indicate that the methodology is identifying modulatory effects that are consistent with what we would expect from prior detailed studies on the mesoscale effects of propofol anesthesia, thus supporting the validity of the proposed approach.

## 4. Discussion

### 4.1. Data-driven inference of modulation for understanding non-stationary brain dynamics

Brain dynamics are fundamentally nonstationary, changing as individuals switch between varying tasks, cycle between sleep and wake states, or as pathologies within the brain improve or worsen. Such non-stationarity is mediated, at least in part, by processes of neuromodulation that potentiate or attenuate synaptic connections within and between brain areas. Our goal in this paper was to develop a data-driven, parametric modeling framework for identifying such modulation at whole-brain scales.

Our specific approach was to formulate modulation within a physiologically interpretable RNN construct, where a baseline set of synaptic weights – common to all non-stationary regimes – is scaled by regime-specific modulation. Within this framework are several nontrivial technical challenges. Specifically, because our model is formulated at the level of latent (unobserved) neural populations, we faced the dual-estimation problem of fitting model parameters and estimating state variables at the same time. Compounding this is the need to fit not one connection matrix, but rather a family of such matrices. Since the base connection weights ***W*** are multiplied by the modulations ***Γ***_*i*_ (*i* = 1, 2, …, *m*), this is a fundamentally ill-posed problem. However, we showed that by imposing appropriate priors on the construction of both ***W*** and ***Γ***, the problem can be made tractable, leading to interpretable results.

At a mathematical level, our modeling setup involves, in essence, a matrix factorization problem: there are a number of distinct effective connectivity matrices (***W*** ⊙ ***Γ***_*i*_), which are then factorized into a component common across all regimes (***W***), and a modulatory component which varies discretely based on regime (***Γ***_*i*_). While there are algorithms for the decomposition of a matrix into the Hadamard product of two matrices [40, 41], these algorithms solve a problem which is constructed differently from ours. The algorithms in [40, 41] decompose a given matrix into two or more low-rank matrices. In our problem, we are principally concerned with finding finding a decomposition with a component matrix *common* to all given matrices, as motivated by our specific domain application context. Importantly, we allow our common component (***W***) to be full-rank, rather than decomposing into two low-rank matrices.

As noted, data-driven decomposition of neuromodulatory effects on mesoscale dynamical models has not been widely studied in the computational neural modeling community. Several authors have, however, done work with a similar motivation for taking a neuromodulation approach to modeling. Li et al [21] developed a statistical generalized linear model (GLM) construct that embodies a modulatory nonstationary architecture for neuronal-level modeling. They specifically imposed a level of similarity between switched models by incorporating a Gaussian prior onto the weight matrices of their GLMs. They further incorporated a decomposition of connection strength from connection direction – decomposing their weight matrices into a direction component in {−1, 0, 1}, and a strength component in ℝ_*>*0_. Having done this decomposition, they also impose a Gumbel-Softmax prior on the direction component, minimizing the number of connections that switch direction with different regimes. In contrast, working at the meso-scale, we enforce a direction mask onto our effective connectivity matrices which does not change over time, and decompose the strength of each matrix into a common component ***W*** and a switched component ***Γ***.

There are certainly many biophysical models that have engaged neuromodulation from a bottom-up perspective, especially at neuronal and small-circuit scales, e.g., [42, 43, 44], which can generate inferences and predictions at the meso- and macro-scale [45, 46]. We view our contribution here as a methodological enabling of such approaches in the sense that our framework takes mesoscale data and performs an inference problem to arrive at a mesoscale model of neural modulation.

### 4.2. Modulated MINDy provides individualized inference of modulated whole-brain dynamics

Our developed approach represents a generalization of our mesoscopic individualized neural dynamics (MINDy) framework [8, 9]. Specifically, rather than identifying a single dynamical regime, the generalized framework proposed here allows for spatially relevant representations of both the base connectivity common to all regimes as well as modulation that may vary with time and brain state. In this sense, this modulated model architecture is also multi-timescale, as the population activity state ***x*** changes on a much faster timescale than the selection index *i*. Importantly, our model architecture is biologically interpretable, providing insight into excitatory-inhibitory dynamics not provided by vanilla RNN architectures. Modulated MINDy accurately estimates known latent dynamics from synthetic data, and infers reliable and individualized models when tested on human data. Modulated MINDy is also scalable, requiring only ~10 minutes to fit 30 minutes of 20-channel EEG data with 3 modulation matrices.

### 4.3. Modulated MINDy as a tool to infer interpretable neuromodulation

As a proof of concept, we tested modulated MINDy on open-source, labeled EEG recordings of subjects receiving the general anesthetic drug propofol. We found that our models had modulation structures consistent with prior literature on propofol sedation, including promoting inhibition along the posterior-anterior spatial axis. Thus, modulated MINDy provides spatiotemporal models which are not only accurate and reliable, but can be interpreted to gain mechanistic insight into an individual’s brain dynamics and how they are modulated over time or as a function of exogenous factors or inputs. We emphasize that our applicative example in anesthesia was not intended to make a specific scientific point about propofol *per se*, but rather as a methdological validity test.

We envision a number of applicative contexts for the proposed method, for both basic scientific and clinical questions. Modulated MINDy can be used in a task context, to infer modulations unique to specific cognitive functions. It could be used, as in this proof of concept, to characterize the spatiotemporal effects of an exogenous intervention, either pharmacological or otherwise. Additionally, it could be used in clinical contexts, to associate different modulation patterns with various states of pathology, e.g., seizures, coma, or ischemia. Ultimately, modulated MINDy is a modeling tool to infer changes that underlie non-stationary brain recordings, where the non-stationary regimes are labeled in the data.

### 4.4. Limitations

We note a few important limitations of the proposed approach. Most notably, as outlined in our introduction, our approach tackles the problem of inferring what is modulated within network dynamics, and not the companion problem of inferring when such modulation has occurred. In other words, we require here known demarcation or labeling of nonstationary regimes. Future work will engage the challenge of inferring both modulation and regime engagement, simultaneously. Our model itself is formulated at the mesoscale, with clear abstraction of cellular and sub-cellular dynamics. These assumptions could in principle be generalized, though that would lead to increasingly computational complexity regarding the ensuing inference problem.

### 4.5. Conclusion

In conclusion, we have presented modulated MINDy, a framework for fitting multiscale, modulated, mesoscale models of brain dynamics to individual data, and have validated it on both ground truth synthetic and actual human EEG data. In the future, we plan to extend this model by enabling modulated inference on data where the modulatory states are unlabeled, i.e., where there is a need to infer also the points at which the non-stationary regimes within the data change. We anticipate that modulated MINDy’s ability to give mechanistic inference will make it a powerful tool for analysis in many neuroscientific and clinical contexts.

## Acknowledgments

Portions of this work were supported by grants R01NS130693 and 5T32NS126157-02 from the US National Institutes of Health.

